# Modeling Neuromotor Adaptation to Pulsed Torque Assistance During Walking

**DOI:** 10.1101/2024.02.19.580556

**Authors:** GilHwan Kim, Fabrizio Sergi

**Affiliations:** Department of Mechanical Engineering, University of Delaware, Newark, DE 19716, USA; Department of Biomedical Engineering, University of Delaware, Newark DE, 19713, USA

## Abstract

Multiple mechanisms of motor learning contribute to the response of individuals to robot-aided gait training, including error-based learning and use-dependent learning. Previous models described either of these mechanisms, but not both, and their relevance to gait training is unknown. In this paper, we establish the validity of existing models to describe the response of healthy individuals to robot-aided training of propulsion via a robotic exoskeleton, and propose a new model that accounts for both use-dependent and error-based learning.

We formulated five state-space models to describe the stride-by-stride evolution of metrics of propulsion mechanics during and after robot-assisted training, applied by a hip/knee robotic exoskeleton for 200 consecutive strides. The five models included a single-state, a two-state, a two-state fast and slow, a use-dependent learning (UDL), and a newly-developed modified UDL model, requiring 4, 9, 5, 3, and 4 parameters, respectively. The coefficient of determination (*R*^2^) and Akaike information criterion (AIC) values were calculated to quantify the goodness of fit of each model. Model fit was conducted both at the group and at the individual participant level.

At the group level, the modified UDL model shows the best goodness-of-fit compared to other models in AIC values in 15/16 conditions. At the participant level, both the modified UDL model and the two-state model have significantly better goodness-of-fit compared to the other models. In summary, the modified UDL model is a simple 4-parameter model that achieves similar goodness-of-fit compared to a two-state model requiring 9 parameters. As such, the modified UDL model is a promising model to describe the effects of robot-aided gait training on propulsion mechanics.

## I. INTRODUCTION

Robot-assisted gait training has been implemented in rehabilitation due to an assumed advantage of effectiveness, greater repeatability, and consistency compared to the standard approach [1], [2]. In robot-assisted gait training, the torque/force assistance to be provided during training may be modified based on the specific needs of a participant. By establishing a relationship between the participant response and assistive parameters, we can predict the optimal torque/force profile that yields a desired participant response. To obtain the optimal torque/force assistance profile for an individual, it is important to consider the variability among participants and to balance between the number of observations and the accuracy of modeling the participant’s response [3]. To accomplish both objectives at once, the human-in-the-loop (HIL) optimization method was introduced [4]. HIL was successfully implemented with various optimization methods including gradient descent [5], covariance matrix adaptive evolution strategy (CMA-ES) [6], and Bayesian optimization [7], [8]. Previous research has shown that incorporating additional information about the relationship between control parameters and participant responses boosts optimization [9], primarily because the number of observations required to construct a precise model is reduced. However, previous optimization methods tested in gait training did not account for the participant’s adaptation during gait training as they assumed that participants would yield the same response when the same torque/force intervention was applied, which limits extensions of these method to rehabilitation applications. Therefore, there is a need to develop models of neuromotor adaptation in response to robot-assisted gait training.

Motor adaptation models describe how the internal model used by our central nervous system to control our movements changes in response to changes in task dynamics. To describe how the central nervous system refines these models, error-based learning models are widely used. In an error-based learning model, the internal model is updated by two processes: a feed-forward signal predicted from internal dynamics, and an error-based component that updates the model based on the perceived error. Errors are often calculated based on the difference between the actual and planned movement or force. The two-state error-based learning model is one of motor adaptation models most widely used in previous research that focused on upper-extremity movements [10], [11]. To address use-dependent effects in the human response, where individuals often rely on past movements to decide the current movement, a use-dependent learning (UDL) model was previously introduced [12]. However, the previous UDL model may describe adaptation during training but is not able to address remaining effects after training, which is the ultimate goal of rehabilitation after neurological injury. Also, the previous models have not been validated within the context of walking mechanics.

In this work, we establish whether state-space models of motor learning can describe the effects of torque pulse applied to the hip and knee joint during the stance phase of walking on measurements of propulsion mechanics, using available data previously collected by our lab [13]. Moreover, we formulated a modified use-dependent learning (modified UDL) model by introducing a reference state update equation in the existing UDL model, to account for the possibility of after-effects of gait training. State space models including single-state, UDL model, two-state fast and slow model, two-state model, and modified UDL model were compared in terms of goodness of fit to the participant- and group-level responses during and after gait training. In our analysis, we tested the hypothesis that the modified UDL model better describes participant responses, both during and after gait training, compared to existing models.

## II. Methods

### A. Data Collection and Processing

In a previous research study, sixteen healthy participants were exposed to pulses of torque to the hip and knee joint while walking on the treadmill to modulate propulsion mechanics [13] (Fig. 1a). Eight different torque pulse patterns used in this research were generated based on the combination of pulse timing and amount of pulse torque to the hip and knee joint (Fig. 1b). In each trial, each participant was exposed to the same torque pulse intervention pattern continuously for 200 strides. 100 strides of baseline without torque pulse were conducted before and after torque intervention session. Two outcome measurements were collected: hip extension angle (HE) at peak anterior ground reaction force and normalized propulsive impulse (NPI).

**Fig. 1:**
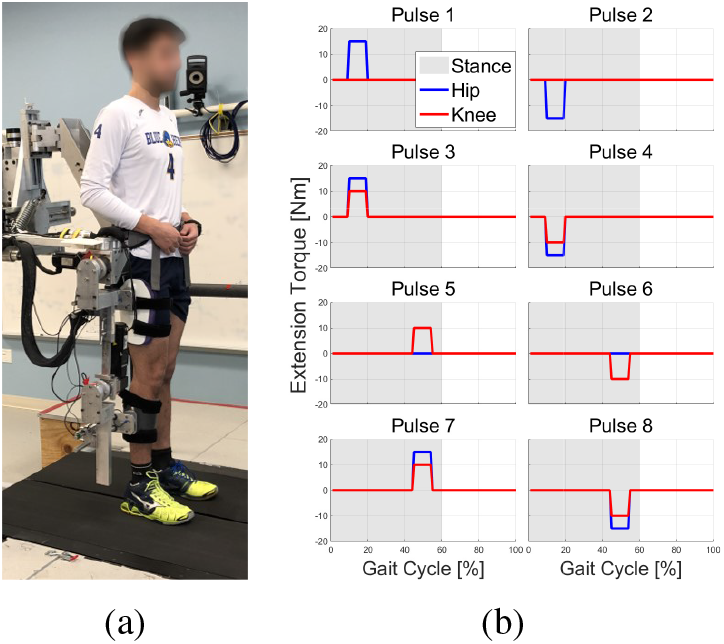
(a) Participant with the Active Leg Exoskeleton II (ALEX II) standing on the instrumented split-belt treadmill. (b) The torque pulse combinations applied to participants.

Outcome measurements were processed separately for each participant, pulse condition, and outcome measurement. Measurements were processed by filling missing data through linear interpolation. To reduce the impact of noise in HE and NPI, the response at each stride was averaged with two preceding and two subsequent responses, separately for each session: baseline, torque intervention, and after-effect. Given that participants were not expected to fully adapt to the experiment setting during the early baseline, responses from the 81st to the 100th strides in the baseline were averaged and subtracted from the overall responses to establish a reference response of zero value. The overall responses were normalized by dividing them by the standard deviation of responses from 81th to the end of strides of each experiment.

### B. Modeling Use-Dependent Learning in the Response to Pulsed Torque Assistance

Five different motor adaptation models were used to describe participant responses data from Sec. II-A. To describe gradual and non-exponential changes of participant response during and early after training, as well as the positive association between effects of training and after-effects of training previous observed in our lab [13], we selected the UDL model [12]. However, because the original UDL model could not account for asymptotic after-effects, we modified the UDL model by introducing a reference state update equation. In addition to these UDL models, standard models of neuromotor adaptation were also included. Specifically, a two-state fast and slow model was selected since this model was often used in previous upper extremity research describing adaptation behavior during and right after training [10], [11]. For a more extensive comparison, a simpler version of this model (single-state model), and a general formulation of a two-state model were also included, as described below.

#### 1) Single-State Model

The single-state model describes error updates and retention of the previously planned movement in a simple way. The movement planned at next stride *x*(*n* + 1) is defined as a function of the previously planned movement *x*(*n*) and error signal *e*(*n*). In our work, *x* refers to the planned hip extension angle (HE) or normalized propulsive impulse (NPI), while *u* is the input applied by the exoskeleton, which induces adaptations in propulsion mechanics. The model equations are as follows:

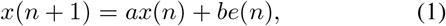

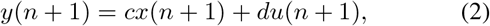

where *a* is the retention coefficient (it indicates how the previously planned movement affects the next movement), *b* is the error update coefficient (it indicates how the previously experienced error affects the next movement), and *c* and *d* indicate how the observed output *y* (i.e., HE or NPI in our data) is affected by the movement plan and by the applied perturbation *u*. Error *e*(*n*) is calculated as the difference between the planned and actual movement, i.e., *e*(*n*) = *y*(*n*) *−x*(*n*). The equation above can be transformed in the standard discrete-time state-space form as

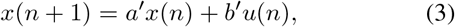

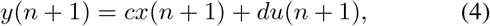

where *a*^*′*^ = *a* − *b* + *bc, b*^*′*^ = *bd*, as a 4-parameter model (*a*^*′*^, *b*^*′*^, *c*, and *d*).

#### 2) Two-State Fast and Slow Model

In this model, a movement plan *x*(*n*) is defined from the linear combination of two separate states: a fast and a slow state [10]. Each state has different dynamics in that the fast state reacts to a perturbation and returns to a reference value faster than the slow state. Since the slow state contains error feedback effects induced from the perturbation, the system may exhibit temporary after-effects when training is completed. The model equations are expressed as:

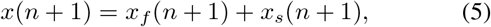

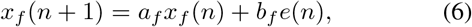

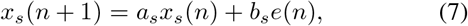

where *e*(*n*) is described as error between planned movement *x*(*n*) and observed output *y*(*n*). Since the slow state (*x*_*s*_) react slower than fast state (*x*_*f*_), the retention and error update coefficients are constrained to verify the following equations: *a*_*s*_ *> a*_*f*_ and *b*_*f*_ *> b*_*s*_. This model can be converted in discrete-time state space form as:

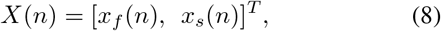

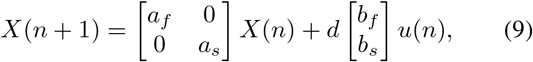

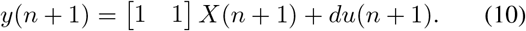

The two-state fast and slow model is defined via 5 parameters (*a*_*f*_, *b*_*f*_, *a*_*s*_, *b*_*s*_, and *d*). When coefficient *d* is equal to zero, the model becomes identical as described in previous implementation in upper extremity experiments [10], [11].

#### 3) Two-State Model

The two-state fast and slow model is an instance of a two-state model of motor adaptation. Using the same error definition, procedure shown in Sec. II-B.1, a two-state model is expressed in the most general form as:

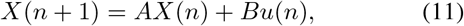

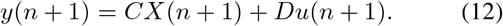

The model is defined via 9 parameters, which are the component of state matrices *A, B, C*, and *D*.

#### 4) Use-dependent Learning Model

The use-dependent learning (UDL) model is a single-state of motor adaptation. The UDL model describes the use-dependent aspect of the human response observed during perturbation. This mechanism is observed simultaneously with error based learning and prediction of movement [12]. The model equation is:

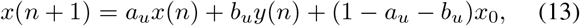

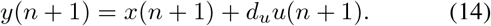

Parameter *a*_*u*_ accounts for retention, while *b*_*u*_ accounts for use-dependent learning, i.e., how the previous output affects the next planned movement. In this model, *x*_0_ is the reference movement in absence of any perturbation, and it is assumed to be a constant. Thus *x*_0_ would account for the reference movement before, during, and after training. As the response data in our experiment results were processed to set the reference movements to zero, *x*_0_ was set to be zero. Therefore, Eq. 13 can be transformed in standard state-space form as:

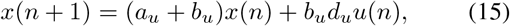

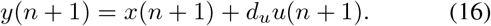

The UDL model is defined via 3 parameters (*a*_*u*_, *b*_*u*_, and *d*_*u*_).

#### 5) Modified Use-dependent Learning Model

The modified use-dependent learning (modified UDL) model is a two-state model describing error-based and use-dependent learning occurring during motor adaptation. The modified UDL model is derived by modifying the existing UDL model, introducing an update equation for the reference state, which will allow the model to describe residual effects induced by training. The model equation is

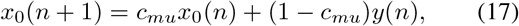

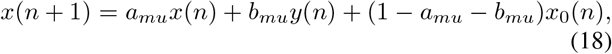

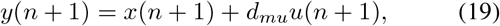

*c*_*mu*_ is the reference state retention parameter, bounded on [0 1]. *a*_*mu*_ is the previous movement retention parameter, and *b*_*mu*_ is the use-dependent learning term. The standard state-space form for the modified UDL model with state vector *X*(*n*) = [*x*_0_(*n*), *x*(*n*)]^*T*^ is

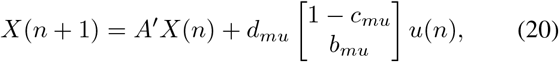

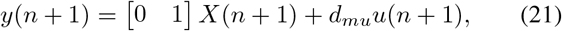

where 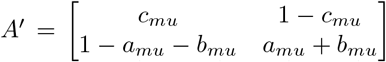. 4 parameters *a*_*mu*_, *b*_*mu*_, *c*_*mu*_, and *d*_*mu*_) define the modified UDL model.

All constraints and number of parameter to be estimated for each motor adaptation model are summarized in Table I.

**TABLE I:** Model parameters and constraints.

| Motor adaptation model | # of parameter | Constraints |
| --- | --- | --- |
| Single-state | 4 | Stability |
| UDL | 3 | Stability |
| Two-state Fast and slow | 5 | $a_s > a_f > 0$,<br>$b_f > b_s$,<br>Stability |
| Modified UDL | 4 | $c_{mu} \in [0 \ 1]$,<br>Stability |
| Two-state | 9 | Stability |

### C. Motor Adaptation Model Fitting

The five motor adaptation models from Sec. II-B were used to describe the data collected as described in Sec. II-A. Group-averaged and individual participant response data were provided for each torque pulse condition for both HE and NPI. Responses from the 81st stride to the end of the experiment were used to fit each motor adaptation model for each torque pulse condition. Constrained state-space model estimation was conducted using MATLAB (MathWorks, Natick, MA, USA) to find optimal values of coefficients parameters in each motor adaptation model equations described in Sec. II-B in the least squares sense. With given initial parameter values, optimal parameter values were searched using the interior-point method [14] or sequential quadratic programming [15]. Model fits were constrained to achieve a stable response, and to satisfy the additional validity constraints listed in Table I. To avoid local minima, fitting was repeated 100 times with different initial parameter values for the two-state fast and slow, UDL, and modified UDL models. Initial values for the single- and two-state models were determined based on the results of UDL and modified UDL models, respectively.

The coefficient of determination (*R*^2^) and Akaike information criterion (AIC) [16] were calculated to quantify the goodness-of-fit and the understand the trade-off between model complexity and goodness-of-fit for the set of models considered. The same procedure was conducted for fitting motor adaptation models in individual participant responses.

### D. Statistical Analysis

*R*^2^ and AIC values from all motor adaptation models were analyzed through statistical software JMP (SAS Institute, Cary, NC, USA). The Shapiro-Wilk test was conducted to check the normality of the outcome measures from each motor adaptation models. If the outcome measures were normally distributed, a one-way repeated measures ANOVA test was conducted to check for the existence of statistical difference between models. When there was statistical significance, a paired t-test was conducted to compare the goodness-of-fit between motor adaptation models. Otherwise, when outcome measures were not normally distributed, Friedman’s and Wilcoxon signed rank test were conducted to compare *R*^2^ and AIC values from different models. To correct inference for multiple comparisons, the paired tests were conducted at a false positive rate of *α <* 0.05*/*10, using a Bonferroni correction for 5 motor adaptation models (10 pairs). Comparison between results of two motor adaptation models was conducted for each measurement condition: HE, NPI, both HE and NPI.

## III. Results

### A. Model Fitting to the Group-averaged Response

Four examples of motor adaptation model fitting to the group-averaged response of HE and NPI are shown in Fig. 2. Fitness values are presented in Table II, with the best values of goodness-of-fit highlighted in bold text.

**Fig. 2:**
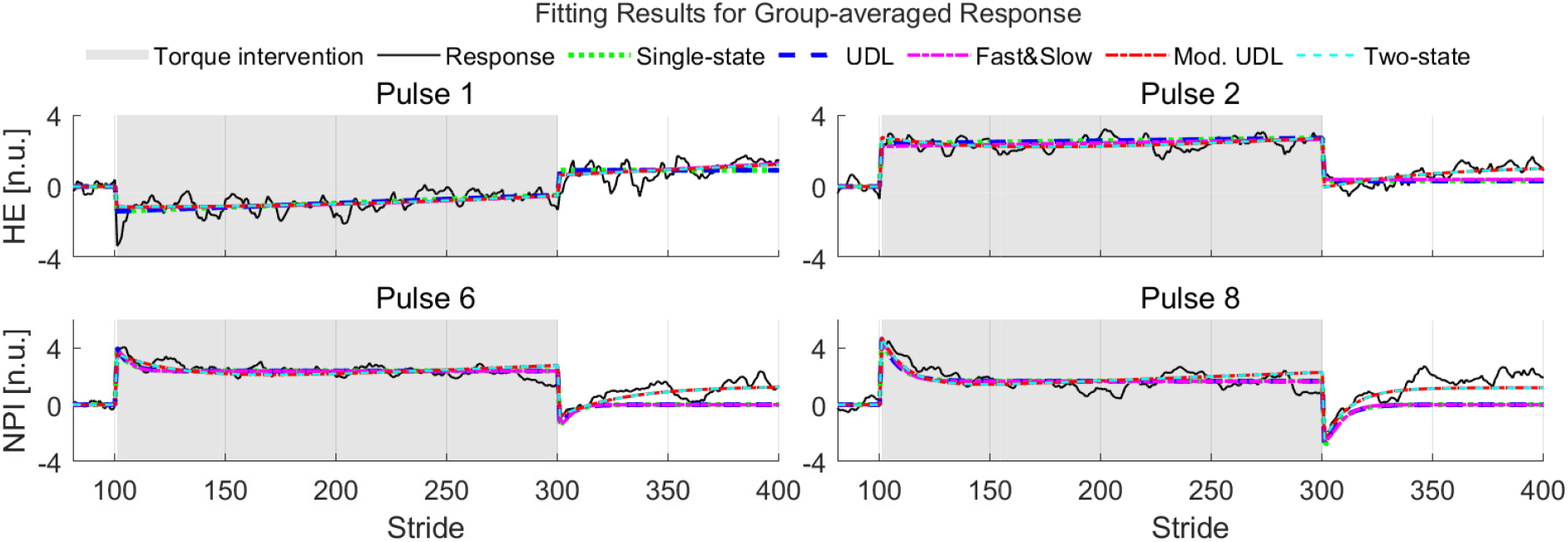
Model fitting for group-averaged responses in hip extension (HE) and normalized propulsive impulse (NPI) during experiments with selected torque pulse conditions. The gray area corresponds to the torque intervention session, where participants experienced torque pulses while walking on the treadmill. Corresponding fitness values (*R*^2^ and AIC) are denoted in Table II.

**TABLE II:** Fitness values for group-averaged HE (NPI)

| Pulse condition |  | Pulse 1 | Pulse 2 | Pulse 3 | Pulse 4 |
| --- | --- | --- | --- | --- | --- |
| Single-state | $R^2$ | 0.76<br>(0.34) | 0.78<br>(0.74) | 0.80<br>(0.38) | 0.94<br>(0.47) |
|  | AIC | 464<br>(785) | 438<br>(481) | 395<br>(761) | -0.2<br>(715) |
| UDL | $R^2$ | 0.76<br>(0.34) | 0.78<br>(0.74) | 0.80<br>(0.38) | 0.94<br>(0.47) |
|  | AIC | 462<br>(783) | 436<br>(479) | 393<br>(759) | -2.2<br>(713) |
| Fast&Slow | $R^2$ | **0.77**<br>(0.32) | 0.80<br>(0.75) | **0.81**<br>(**0.45**) | **0.95**<br>(0.48) |
|  | AIC | 445<br>(793) | 395<br>(475) | 392<br>(727) | -22<br>(706) |
| Mod. UDL | $R^2$ | **0.77**<br>(**0.41**) | **0.88**<br>(**0.77**) | 0.81<br>(0.45) | **0.95**<br>(**0.52**) |
|  | AIC | **443**<br>(**746**) | **245**<br>(**443**) | **385**<br>(**726**) | **-24**<br>(**678**) |
| Two-state | $R^2$ | **0.77**<br>(**0.41**) | **0.88**<br>(**0.77**) | **0.81**<br>(0.45) | **0.95**<br>(**0.52**) |
|  | AIC | 453<br>(756) | 255<br>(453) | 395<br>(736) | -14<br>(688) |
| Pulse condition |  | Pulse 5 | Pulse 6 | Pulse 7 | Pulse 8 |
| Single-state | $R^2$ | 0.96<br>(0.78) | 0.92<br>(0.49) | 0.98<br>(0.73) | 0.99<br>(-0.05) |
|  | AIC | -121<br>(426) | 98<br>(702) | -347<br>(501) | -441<br>(930) |
| UDL | $R^2$ | 0.96<br>(0.78) | 0.92<br>(0.49) | 0.98<br>(0.73) | 0.99<br>(-0.05) |
|  | AIC | -123<br>(424) | 96<br>(700) | -349<br>(499) | -443<br>(928) |
| Fast&Slow | $R^2$ | 0.96<br>(**0.79**) | 0.92<br>(0.49) | 0.98<br>(0.74) | 0.99<br>(-0.04) |
|  | AIC | -116<br>(418) | 93<br>(703) | -345<br>(492) | -467<br>(931) |
| Mod. UDL | $R^2$ | **0.97**<br>(**0.79**) | **0.93**<br>(**0.77**) | 0.98<br>(0.73) | **0.99**<br>(**0.57**) |
|  | AIC | **-164**<br>(**416**) | **77**<br>(**447**) | **-383**<br>(501) | **-558**<br>(**643**) |
| Two-state | $R^2$ | **0.97**<br>(**0.79**) | **0.93**<br>(**0.77**) | **0.98**<br>(**0.74**) | **0.99**<br>(**0.57**) |
|  | AIC | -154<br>(426) | 87<br>(457) | -378<br>(**489**) | -548<br>(653) |

UDL, modified UDL < two-state fast and slow, two-state < single-state, and modified UDL < two-state model, for outcome measurement of HE. There was no difference group for the outcome measurement of NPI. For both HE and NPI, except for three pairs (single-state vs. two-state fast and slow, UDL vs. two-state fast and slow, and two-state fast and slow vs. two-state model), all other pairs showed significant difference at the corrected false positive ratio of 0.005.

With the exception of *R*^2^ values obtained from the single-state, UDL, and two-state fast and slow model, all other values, including AIC and *R*^2^ values, exhibited non-normal distributions. The Friedman’s test results showed that there was a significant effect of model on *R*^2^ and AIC values for all outcomes (HE, NPI) with *p*-values smaller than 0.001. Post-hoc Wilcoxon signed rank test also showed significant difference in *R*^2^ values among five model pairs: modified UDL *>* single-state, modified UDL *>* UDL, two-state *>* single-state, two-state *>* UDL, and two-state *>* modified UDL, for outcome measurement of HE. These comparisons were significant at the corrected false positive ratio of 0.005. For outcome measure of NPI, two pairs showed significant difference: two-state *>* single-state, and two-state *>* UDL. When considering both HE and NPI, all pairs showed statistical difference in *R*^2^ values. For the Wilcoxon signed rank test for AIC values, five pairs showed significant differences: modified UDL *<* single-state, modified UDL *<* UDL, modified UDL *<* two-state fast and slow, two-state *<* single-state, and modified UDL *<* two-state model, for outcome measurement of HE. There was no difference group for the outcome measurement of NPI. For both HE and NPI, except for three pairs (single-state vs. two-state fast and slow, UDL vs. two-state fast and slow, and two-state fast and slow vs. two-state model), all other pairs showed significant difference at the corrected false positive ratio of 0.005.

### B. Model Fitting to the Participant-level Response

All fitness values, including *R*^2^ and AIC, from five motor adaptation models showed normal distribution. For both *R*^2^ and AIC values, one-way repeated measures ANOVA results showed that there exists a significantly effect of model on goodness of fit for all outcome measurement cases (HE, NPI, and both HE and NPI) with *p*-values smaller than 0.001.

*R*^2^ values showed significant difference for all pairs made from the 5 different motor adaptation models, except for the pair of single-state vs. UDL model. The resulting ranking of *R*^2^ values, from smallest to greatest, was: UDL, single-state, two-state fast and slow, modified UDL, two-state. Statistical significance was maintained in all outcome measurement cases: HE, NPI, and both HE and NPI.

AIC values for fitting HE responses of participants indicated that all pairs of models were significantly different, except modified UDL vs. two-state model. The resulting ranking of AIC values, from greatest (worse fit) to smallest, was: single-state, UDL, two-state fast and slow, modified UDL, two-state. For NPI, UDL vs. two-state fast and slow model showed no statistical difference. For all response of HE and NPI, all pairs showed statistical difference, with the same ranking reported above for HE. Overall, the two-state model and the modified UDL model showed significantly better AIC values compared to other motor adaptation models.

In all cases of participant-level response fitting, the modified UDL model and the two-state model demonstrated better performance compared to the other models. *R*^2^ values indicated that the two-state model showed significantly better fitness performance compared to the modified UDL model (Fig. 3).

**Fig. 3:**
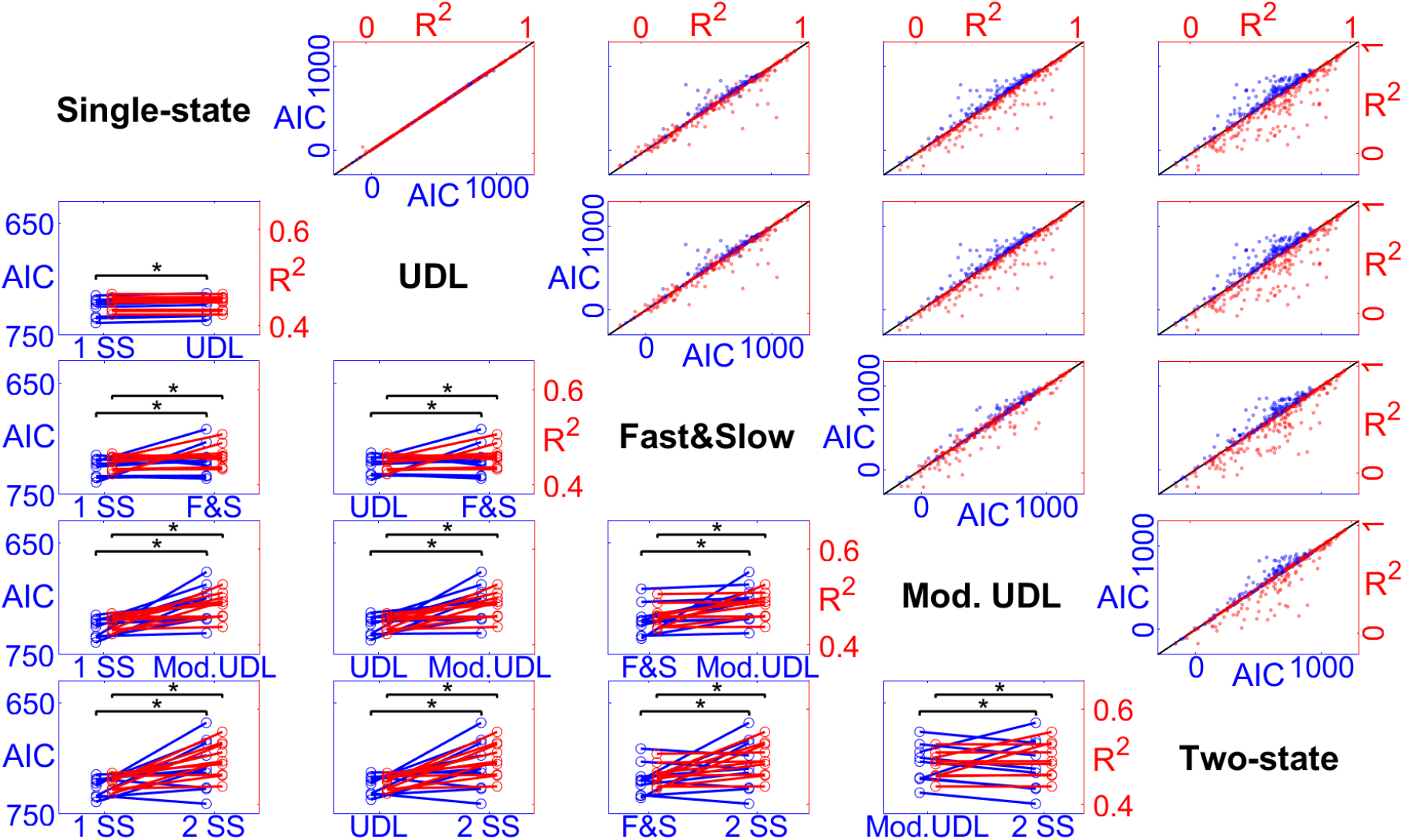
Paired comparisons of *R*^2^ and AIC, reported as scatter plots (top-right), and paired line plots (bottom-left). Each row/column corresponds to one of the five models, and paired comparisons between models are reported in the intersecting cell. For the scatter plots, the unity line is provided in black, with *R*^2^ values provided as red dots, and AIC values in blue. Large departures from the unity line are an indication of large differences of goodness-of-fit. Asterisks indicate significant differences at the corrected false-positive rate. For the paired line plots, randomly undersampled ten outcomes between AIC values of 650 to 750 and *R*^2^ values of 0.4 to 0.6 from all outcome measurement cases (HE and NPI) are displayed for improved figure clarity. Also, the AIC scale is inverted to align with the changes in *R*^2^ values between different models.

## VI. Discussion and Conclusions

In this work, we compared the performance of five different motor adaptation models describing group-averaged and individual participant responses collected in a previous experiment [13]. Based on the goodness-of-fit values (*R*^2^ and AIC), the modified UDL model is expected to be the best model for describing group-averaged responses. For addressing participant-level responses, the modified UDL model and the two-state model are similarly suitable for describing the response of HE, and the two-state model has shown to be superior for describing the response of NPI.

In most of cases from participant-level and group-averaged fitting results, motor adaptation models were found to be more accurate to describe response of HE than NPI (*R*^2^ values for HE: 0.61±0.27, NPI: 0.40 ±0.25). Robotic intervention resulted in a greater effect on propulsion kinematics, specifically HE, compared to propulsion kinetics, specifically NPI. This may be due to both a smaller effect introduced by the robot and to the fact that NPI may contains more noise, which was not fully eliminated via data processing procedure. Therefore, in describing kinematic outcome measurements, motor adaptation models may be expected to be more effective than implementing them for kinetic outcome measurements.

Since the group-averaged data contain less noise compared to participant-level responses, this may effect the goodness-of-fit for describing responses using motor adaptation models, as demonstrated by the differences in *R*^2^ values (group-averaged: 0.72*±*0.25, participant-level: 0.49*±*0.28).

In terms of the capability of motor adaptation models to describe remaining after-effects, both the single-state and the UDL model will eventually exhibit no after-effects when a large number of strides for after training session are provided in experiment. Therefore, for further implementation in gait training targeting after-effect modulation, the modified UDL and the two-state models should be considered. When considering fitness values to describe both HE and NPI from group-level responses, the modified UDL model performed better than the two-state model. However, this relationship was reversed when addressing participant-level response of NPI. This result suggests the possibility that the two-state model performs better with noisy responses, while the modified UDL model may be superior in absence of noise. The state-space model with two states can be transformed into a discrete transfer function to calculate the current output based on 5 values: the current and the two past inputs, and two past responses. As such, a five-parameter model would be sufficient to describe the two-state model, without explicitly modeling the structure of process and measurement noise. Therefore, the performance difference between the modified UDL and the two-state model describing group- and subject-level responses were mainly due to the structure constraints of state matrices in modified UDL model, and to the effect of modeling noise propagation in these two different formulations. For further research targeting kinematic variables such as HE or trailing limb angle during and after gait training using HIL optimization method, the modified UDL model is expected to perform best with the additional noise elimination procedure during gait training. With proper calibration of the prediction accuracy, the modified UDL model may be used to implement model-based predictions of after-effects of training.

